# MultiSpecies Canopy Segmentation: Interactive Machine-Learning and Pseudo-Labelling are key

**DOI:** 10.64898/2025.12.07.692840

**Authors:** Charles Rongione, Abraham George Smith, Céline Chevalier, Christophe De Vleeschouwer, Guillaume Lobet, Xavier Draye

## Abstract

This study investigates the challenge of creating datasets for training multiclass deep-learning segmentation models, specifically for segmenting multi-species canopy images. Creating training sets for deep-learning based segmentation of multispecies canopies is currently too labor-intensive and time-consuming to be viable. To address this challenge, we propose a novel pipeline that uses fully convolutional neural networks (FCNNs) to transition from single-species images to segmented multi-species images. This paper demonstrates that FCNNs can effectively generalize learning from single-species canopy images to multispecies canopy images, achieving accurate pixel classification in mixed species canopies even when the network was trained only on images of single-species canopies. Additionally, we introduce Interactive Machine Learning and pseudo labeling as a method for generating a single-species canopy training set in a matter of minutes. We also present two software packages to implement our approach and extensively evaluate them against several baselines. Our findings demonstrate that our approach can significantly reduce the human time load required for semantic segmentation of multispecies canopy images, achieving over 90% accuracy in less than 10 minutes. This new method has the potential to greatly facilitate the study of multispecies canopies.

## 1 Introduction

### 1.1 Context

Sustainable intensification is the practice of increasing food production in an environmentally and socially responsible way. Mixed cropping is an approach that involves planting multiple crops in the same field. This practice has several benefits, including reduced use of inputs like water and fertilizers Guvenc2008, increased crop diversity and resilience Iqbal2019, and improved soil health tang2020cassava. Mixed cropping allows farmers to achieve multiple goals simultaneously, maintaining their yields while protecting natural resources. One challenge in the practical use of mixed cropping is the need to carefully select and combine crops to minimize competition and maximize compatibility and complementarity. The state of knowledge about designing efficient mixed cropping systems is currently insufficient due to their complexity and the limitations of current phenotyping methods used to quantify such complexity. High throughput phenotyping is a promising approach for characterizing the complexity of mixed cropping systems and for understanding their potential benefits and challenges. However, this approach is not yet sufficiently developed for the study of mixed cropping systems.

Deep learning based image segmentation is a promising approach for high throughput phenotyping of mixed cropping systems through the analysis of their canopies. Image segmentation is the process of dividing an image into distinct regions or segments, and deep learning algorithms can be used to automatically identify and classify different plant species in an image. This approach has several potential benefits for the study of mixed cropping systems. First, it allows for the rapid and automated measurement of multiple plant traits, including leaf area, canopy structure, and species composition. Second, it can be used to study mixed cropping systems at different scales, from small plot trials to larger field-scale experiments. This approach has the potential to ease the process of quantifying these complex systems and, therefore, to better inform the development of more sustainable and efficient mixed cropping systems.

One of the main limitations of deep learning image segmentation is the need to create a large and diverse training set in order to accurately train the algorithm. A training set is a collection of images and corresponding labels that are used to train the algorithm to recognize and classify different objects or classes in an image. Creating such a training set can be time-consuming and resource-intensive, as it currently requires to manually label a large number of images with the appropriate class labels. It is also important to ensure that the training set is diverse and representative of the range of conditions and classes that the algorithm will encounter in real-world applications. Because Mixed cropping systems can involve a wide range of plant species, canopy structures, and environmental conditions, it is difficult to create a training set that accurately represents the range of classes and the complexity that the algorithm will encounter.

### 1.2 This work

The purpose of this study is to investigate potential methods for reducing the time and effort required for manual annotation in the use of fully convolutional neural networks for segmenting multi-species canopy images. To achieve this goal, we focus on addressing a key challenge of supervised segmentation: obtaining sufficient quantities of training data in the case of multispecies canopies.

To that end, we propose a five step pipeline for transitioning from single-species images to segmented multi-species images (see figure 1):

**Figure 1.**
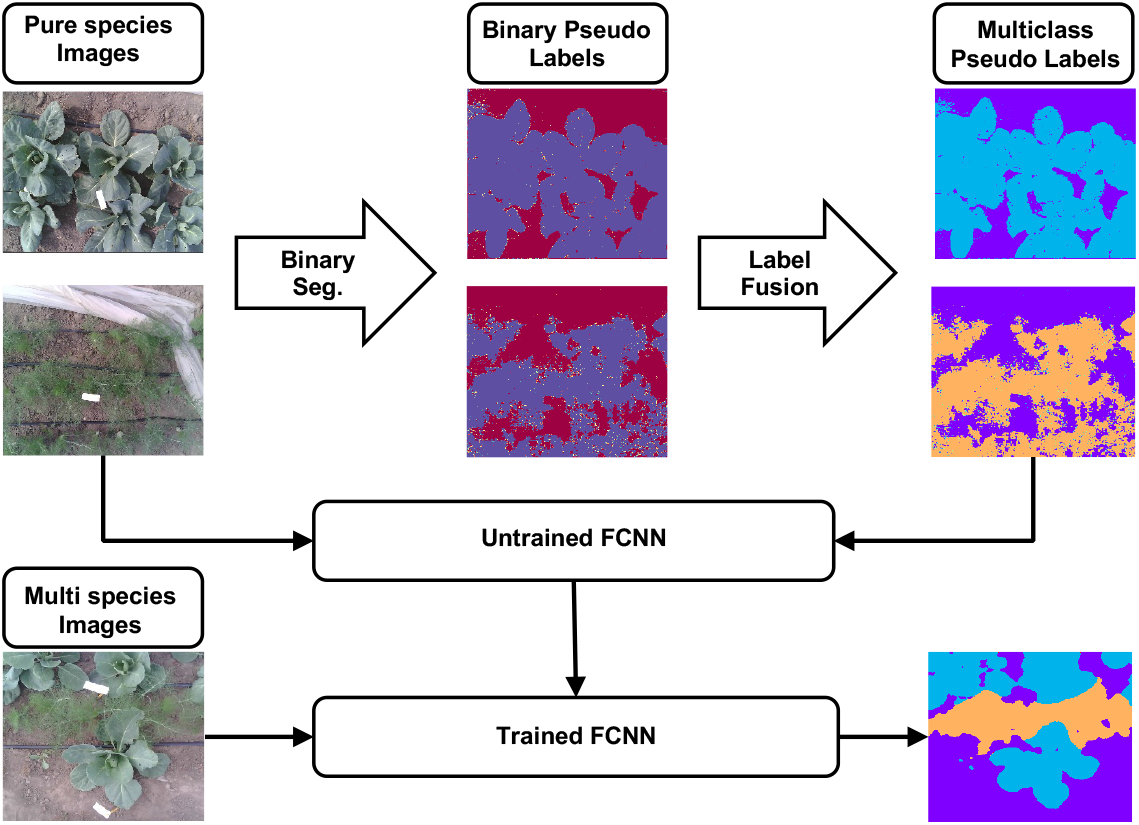
Schematic view of the workflow of the segmentation pipeline, starting with mono-species canopies interactive segmentation to generate binary-pseudo labels, use of their species names to create multi class pseudo-labels (fennel in orange, cabbage in blue) and the final inference on multi-species canopies.

1. Collection of images of pure crops with the species indicated
2. Generation of binary pseudo-labels for these images with Interactive Machine Learning
3. Conversion of these binary pseudo-labels into multi-class pseudo-labels using species indications of the images
4. Training of a fully convolutional neural network on these images and their multi-class pseudo-labels obtained in the previous step
5. Use of the trained fully convolutional neural network to predict segmentation masks on images of multispecies canopies

To perform the second step, we adopt an Interactive Machine Learning (IML) approach, i.e., using a combination of human input and machine learning algorithms to label images. This method is more efficient than traditional manual annotation techniques because only a fraction of the pixels of an image needs to be manually annotated. Based on those manual annotations, the model will annotate the rest of the image itself. Those model-generated annotations are called pseudo-labels (PL), in constrast with manual annotation. We then use those interactively generated PL to train a fully convolutional neural network to generate PL on the remaining images of the dataset, without the need of additional human intervention. As part of step 3, those binary PL will then be classified according to the species of the plant and become multiclass PL. Those multiclass PL constitute the final multiclass training/validation sets.

Because such a combination of IML and pseudo-labeling are not well studied in the literature, we compare multiple protocols that combine IML and pseudo-labels to evaluate their effectiveness in creating training sets for our task. We present two software tools, LeafPainter and RootPainter, as options for implementing these protocols. LeafPainter is designed for use on low-resource computers, while RootPainter is suitable for more powerful setups with GPUs. Through this comparison, we aim to identify the most effective and efficient method for generating labels for deep learning-based semantic segmentation of mixed crop images.

Our results demonstrate that both RootPainter and LeafPainter approaches outperform all of the compared baselines, including the widely used Ilastik software, in terms of segmentation accuracy on our multispecies canopies dataset. Not only does this approach achieve high levels of accuracy (over 90%), but it also greatly reduces the amount of labor required for semantic segmentation. With only a few minutes of human input, our approach offers an efficient solution for studying multispecies canopies.

The originality of our work is:

- Development of LeafPainter, a gradient boosted decision trees-based software for generating pseudo-labels for a segmentation task
- Evaluation of RootPainter on a canopy segmentation task
- Comparison on several pseudo-labels generation protocols based on Interactive Machine Learning
- Use of mono-species images to segment multi-species canopies images

### 1.3 State of the art: Deep learning based image segmentation for canopies analysis

Fully convolutional neural networks have emerged as the state-of-the-art method for image segmentation in recent years, and have been widely applied and validated in plant image analysis (for example, Jiang2020; kamilaris2018; Santos2020;Ubbens2017). FCNNs are able to learn the architectural features of plants, such as leaf shape, stem shape, and colors, automatically, without the need for manual feature identification or creation Tang2017. This is particularly useful in the context of heterogeneous images taken in the field, where manually encoded features may not be robust enough for use in precision agriculture (Lottes2018; McCarthy2010). In fact, it has been shown that the greater the heterogeneity of the images, the more necessary it becomes to use FCNNs for image segmentation (Lottes2018; McCarthy2010). This is particularly relevant for multi-species segmentation, where the scene is naturally very heterogeneous.

One major constraint of using FCNNs for multiclass semantic segmentation is the need for a large amount of annotation work for outdoor images. In order to train FCNNs for this task, it is necessary to provide examples of the task we want the model to perform, which involves annotating crop images by drawing areas containing plants and assigning classes to these areas corresponding to their species. This can be a time-consuming and costly process, with some studies reporting that it can take up to 10 hours of work per image FuentesPacheco2019. In cases where species are highly intricate, the areas to be annotated can be difficult to label accurately. This requirement for extensive annotation can be a major barrier to using FCNNs for multiclass semantic segmentation of outdoor images. More and more authors are trying to overcome this constraint. We see, for example, the emergence of the use of synthetic datasets (Barth2019; Picon2022).

We are also seeing a resurgence of Interactive Machine Learning. It is a type of machine learning that involves human interaction in the training and labeling process to guide the machine learning model. It can improve the accuracy and efficiency of machine learning models, particularly when working with large datasets dudley2018review. IML involves methods such as manually labeling data points, providing feedback on model predictions, or adjusting model parameters. It leverages the strengths of both humans and machines to improve the performance of machine learning algorithms and achieve better results.

Corrective annotation is a type of Interactive Machine Learning that involves a human providing feedback to a model during the training process. This type of training is commonly used in many domains, including in plant science. Two popular tools that provide a user interface to apply it are RootPainter smith2022RootPainter and Ilastik berg2019ilastik.

RootPainter is a software that utilizes a fully convolutional neural network. During the training process, the user corrects any mistakes made by the model, incrementally expanding the dataset based on these corrections. It has extensively been used for roots and soil analysis (for example seethepalli2021rhizovision, han2021digging, bauer2022development, rinehart2022method) but never for canopies, despite its potential to do so.

Ilastik works the same way, except that it is based on random forests. Between each correction, the model runs a complete training loop, which makes it slower. It has been used for a wider range of problems, including canopies visschers2018quantification.

## 2 Material and methods

This study conducts two empirical evaluations to investigate the efficacy of different binary pseudo-labels generation protocols. Specifically, the following protocols are compared: LeafPainter, Root-Painter, Ilastik, and random patches annotation, with their detailed descriptions provided in subsequent sections. To assess the performance of these protocols, we compare the generated pseudo-labels of 28 images of monospecies canopies with their corresponding manual annotations, using the accuracy metric to quantify similarities based on the percentage of correctly classified pixels. We ensured that our dataset had a balanced distribution to mitigate any potential bias in accuracy due to class imbalance, which is a common concern with this metric. In the second evaluation, we assess the ability of a FCNN to learn multispecies canopy segmentation by training on single species images and their corresponding pseudo-labels generated by the aforementioned protocols. To measure the accuracy of the generated pseudo-labels, we use 28 manually annotated images of multispecies canopies, comparing the segmented output against the ground truth labels. Accuracy is also used as a measure of segmentation quality.

### 2.1 The dataset

The dataset, from which are drawn the previously mentioned test images, consists of 2386 zenithal RGB images of experimental plots, each containing either a single vegetable species or a mixture of two different vegetables. There are four vegetable species in total: fennel, dwarf bean, cabbage, and kale. The dataset includes images of each species in pure culture as well as all possible combinations of two vegetables. Each image is labeled with the name of the species depicted. These labels are important because they allow for the assumption that the plant species of each pixel in the “plant” class is known for pure crop images.

Each plot was photographed four times over the course of the four-month (July, August, September, October 2021) experiment, resulting in images of the plants at different stages of growth and under varying sunlight conditions. The photos were taken in Belgium at either the Lauzelle farm (50.680051034554026, 4.617946520106397) or one of 20 volunteer market gardeners’ plots. The resulting images exhibit a range of contexts, including variations in soil types, crop arrangements, and non-plant objects present on the plots, as well as differences in cultivation techniques among market gardeners. The acquisition of the images was carried out within the framework of Chevalier2022. The cameras used consisted of 2 OEM OV2640 (Omnivision) camera modules, RGB, 2Mpixel, embedded in an ESP32-Cam module (https://docs.ai-thinker.com/en/esp32-cam), installed parallel to each other.

### 2.2 Step 1: collection of images of pure cultures with the species indicated

The mono-species images in this dataset were collected along with the multi-species images (see Chevalier2022 for further details). As these images depict a single species, all pixels classified as “plant” can also be classified as the specific species depicted. The indications of the species present in the images are in the name of the images.

### 2.3 Step 2: Binary pseudo-labeling of the pure culture images

#### 2.3.1 Binary masks generation with LeafPainter

The objective of this step is to conduct binary segmentation on the pure culture plot images, categorizing each pixel as either “plant” or “background.” To accomplish this, we propose an iterative method that aims to annotate only the areas of the images where the predictions are incorrect. This method consists of the following six steps:

1. Annotate few areas of soil and plant and store these annotations in a dataset.
2. Split the dataset into a training and validation set.
3. Train a gradient boosted decision trees model on the training set until performance ceases to improve on the validation set.
4. Make predictions on the remaining portion of the image.
5. Annotate some areas where the model is incorrect and add these annotations to the dataset.
6. Repeat from step 2.

Upon achieving satisfactory segmentation performance, the user can download the generated mask and repeat the process for subsequent images. Through sufficient annotation, the model will eventually become robust enough to segment the remainder of the dataset without further manual input.

To implement this method, we developed an application called LeafPainter, a user interface that allows the user to complete all steps within the same window and then download the generated binary mask. LeafPainter was created using the Python programming language and the Flask library, and the interface was implemented using HTML5, CSS, and JavaScript. A visual demonstration of LeafPainter can be found at https://www.youtube.com/watch?v=lMpzxDt40Lc. The algorithm underlying LeafPainter is an ensemble of gradient boosted decision trees (GBDT), a machine learning technique used for regression and classification tasks. GBDT is an ensemble of weak decision trees, each of which is created in a sequential manner and tasked with correcting the errors of the preceding tree in the sequence breiman1997arcing. We chose to implement GBDT using the LightGBM library version 3.2.0 from Microsoft ke2017lightgbm, which is also the implementation that offers the best performance in terms of speed, which is critical to maintain the interactivity of the algorithm. The most important hyperparameters (according to Boehmke2017) were optimized using a dataset of approximately 10 million pixels, which was generated by annotating randomly chosen areas of 68 images selected to represent each plant species at each growth stage and each month. This diversity ensures that the subset of the dataset is representative of the distribution of the full dataset, and the dataset was also created to have approximately as many plant pixels as backgrounds. Those parameters are listed in table 1

**Table 1:** Parameters of the gradient boodted decision trees of LeafPainter.

| Parameter | Value | Nature of the parameter |
| --- | --- | --- |
| Number of trees | 100 | The number of trees the model will contain at the end of training |
| Learning rate | 0.05 | The constant by which the output of the last tree created will be multiplied before being added to the temporary model. That is, the contribution of each tree to the final output. |
| Trees maximum depth | 8 | The maximum number of floors a tree can have. |
| Minimum samples per leaf | 2 | The minimum amount of samples that can be grouped in a leaf. |

#### 2.3.2 Binary mask generation with RootPainter

For the purpose of training the RootPainter model, images were subdivised into smaller images of dimensions 800x600 pixels. As per the recommendation outlined in smith2022RootPainter, the training dataset was prepared using the corrective annotation protocol explained in smith2022RootPainter. This involved manual annotation of clear examples in the first six images, with the subsequent annotation process being guided by model predictions. The primary objective of this process is to address the issue of false negatives and false positives in the segmentation output. False negatives refer to regions where the plants were missed by the segmentation model, while false positives refer to regions where the model incorrectly identified the background as plant.

#### 2.3.3 Binary mask generation with Ilastik

The Ilastik software was employed to generate pseudo-labels, following the recommended protocol described in berg2019ilastik. The RGB channels of the images were used in this process. It is worth noting that Ilastik operates similarly to LeafPainter, with the key difference being the utilization of random forests instead of gradient boosted decision trees.

#### 2.3.4 Binary mask generation by random patches selection

This last protocols consists in manually annotating random parts of the images without using the feedback of a model. This random annotations are then stored into the training or validation set.

#### 2.3.5 Pseudo-labels generation on the rest of the dataset

Following the generation of binary masks using RootPainter, Leafpainter, Ilastik, or random patches, a fully convolutional neural network is trained using these masks until convergence. The trained network is then used to generate pseudo-labels on the whole dataset, including the manually processed images by the previously mentionned protocols. In order to assess the impact of this additional step, we compared the quality of the binary pseudo labels with and without it. The different protocols were named by adding “With FCNN Training” (WT). when the network was trained and used on unseen images, while “raw” was added when only the pseudo-labels generated on seen images were used. The resulting list of protocols is as follows:

- Random patches selection raw: Random patches of the images were manually annotated.
- Random patches selection (WT): Random patches of the images were manually annotated and used as training data for a FCNN to generate pseudo-labels from the whole dataset.
- Ilastik raw: The Ilastik software was used to generate binary masks from the images.
- Ilastik (WT): The Ilastik software was used to generate binary masks from the images, and these labels were used as training data for a FCNN to generate pseudo-labels from the whole dataset.
- LeafPainter raw: The LeafPainter software was used to generate binary masks from the images.
- LeafPainter (WT): The LeafPainter software was used to generate binary masks from the images, and these labels were used as training data for a FCNN to generate pseudo-labels from the whole dataset.
- RootPainter raw: The RootPainter software was used to generate pseudo-labels from the images.
- RootPainter (WT): The RootPainter software was used to generate binary masks from the images, and these labels were used as training data for a FCNN to generate pseudo-labels from the whole dataset.

To also quantify the time efficiency of these protocols, we measured the quality of the generated pseudo labels with different total time investments of 6.25, 12.5, 25, 50, 75, and 100 minutes for each protocol.

### 2.4 Step 3: Conversion of the binary pseudo-labels into multicategories ones

Training a multiclass fully convolutional neural network requires a dataset with multiclass labels. However, the dataset generated in previous steps only has binary labels: “plant” and “non-plant.” To enable training of a multiclass FCNN, we convert these binary labels into multiclass labels, including background, cabbage, bean, fennel, and kale. The conversion process is simple because the binary pseudo-labeled images are of single species and therefore only contain one plant species. The resulting dataset has multiclass pseudo-labels for pure species images, with 5 classes. This dataset can be used to train a multiclass FCNN that can segment images into multiple classes.

### 2.5 Step 4: Fully convolutional neural network training on the segmented images

Once the multiclass pseudo-labels for the monoculture images were generated, a multiclass FCNN was trained on the segmented images. The training and validation set were respectively 80% and 20% of the dataset. This FCNN was trained to recognize not only the plant pixels, but also the species to which they belong.

#### 2.5.1 Fully convolutional neural network architecture and implementation

We used a fully convolutional neural network architecture with a U-net-like encoder-decoder structure. The encoder was based on a resnet50 architecture He2015, which has been shown to be effective as an encoder for U-net architectures Zhang2018. Instead of using bottom-up convolutions in the decoder, we employed pixel shuffle oversampling, a technique introduced by Shi2016 that involves a series of convolutions that each learn to extrapolate in a specific direction. This method has been demonstrated to be capable of learning more complex and accurate transformations than simple upward convolutions, and has been found to be particularly effective in the domain of superresolution, which involves increasing the spatial resolution of a signal transmitted by the contraction part of the U-net. The FCNN architecture and training loop were implemented using pyTorch v1.8.1 Paszke2017 and the Fastai library (v2.5.3) Howard2020

#### 2.5.2 Fully convolutional neural network loss function

We implemented a combined loss function consisting of the focal loss Lin2017 and the logarithm of the hyperbolic cosine of the Dice loss Jadon2020, a modified version of the traditional Dice loss Sudre2017 that addresses numerical instability issues Jadon2020. As demonstrated by Wang2020, the focal loss is particularly effective when the dataset contains many small objects, while the Dice loss performs well when the background takes up a large portion of the dataset and focuses on the positive class region (in our case, the plant pixels). Wang2020 showed that combining the dice and focal losses significantly improves performance on their dataset. To the best of our knowledge, this work is the first using this combination of losses.

#### 2.5.3 Fully convolutional neural network hyperparameter tuning

The hyper-parameters of the FCNN (Table 2) were manually tuned with comparisons made by comparing accuracy on the validation set.

**Table 2:**
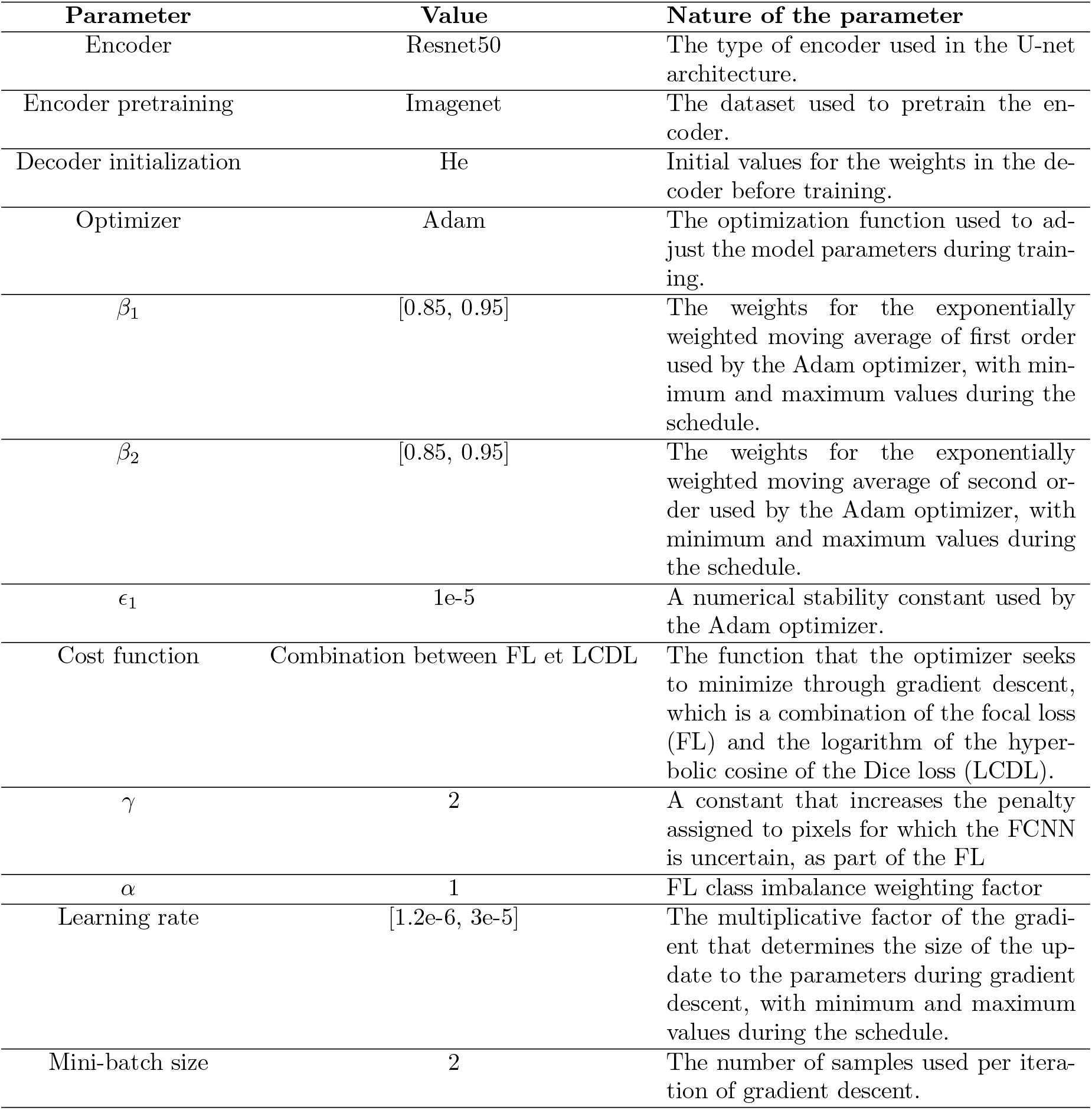
Optimized parameters for the fully convolutional neural network and their nature.

| Parameter | Value | Nature of the parameter |
| --- | --- | --- |
| Encoder | Resnet50 | The type of encoder used in the U-net architecture. |
| Encoder pretraining | Imagenet | The dataset used to pretrain the encoder. |
| Decoder initialization | He | Initial values for the weights in the decoder before training. |
| Optimizer | Adam | The optimization function used to adjust the model parameters during training. |
| $\beta_1$ | [0.85, 0.95] | The weights for the exponentially weighted moving average of first order used by the Adam optimizer, with minimum and maximum values during the schedule. |
| $\beta_2$ | [0.85, 0.95] | The weights for the exponentially weighted moving average of second order used by the Adam optimizer, with minimum and maximum values during the schedule. |
| $\epsilon_1$ | 1e-5 | A numerical stability constant used by the Adam optimizer. |
| Cost function | Combination between FL et LCDL | The function that the optimizer seeks to minimize through gradient descent, which is a combination of the focal loss (FL) and the logarithm of the hyperbolic cosine of the Dice loss (LCDL). |
| $\gamma$ | 2 | A constant that increases the penalty assigned to pixels for which the FCNN is uncertain, as part of the FL |
| $\alpha$ | 1 | FL class imbalance weighting factor |
| Learning rate | [1.2e-6, 3e-5] | The multiplicative factor of the gradient that determines the size of the update to the parameters during gradient descent, with minimum and maximum values during the schedule. |
| Mini-batch size | 2 | The number of samples used per iteration of gradient descent. |

#### 2.5.4 Parameters scheduler

In this study, we used the learning rate scheduler proposed by Smith2019. This scheduler consists of two linear phases: a warm-up phase in which the learning rate increases from the minimum value to the maximum value, and a cooling phase in which the learning rate decreases back to the minimum value. Smith2018 demonstrated that this learning policy should be accompanied by an inverse dynamic for the weights that ponder the exponentially weighted moving average of optimizers using the inertia of the gradient. This means that as the learning rate increases, the weights for the exponentially weighted moving average decrease, and as the learning rate decreases, the weights for the exponentially weighted moving average increase, with a linear relationship between the two. This allows the optimizer to maintain a balance between using the accumulated knowledge from previous iterations (as represented by the exponentially weighted moving average) and adapting to the current gradient.

#### 2.5.5 Data augmentation

Data augmentation is a common technique used in the training of convolutional neural networks. It involves applying transformations to images in order to artificially increase the diversity and richness of the dataset without requiring additional data. Data augmentation can be useful in cases where the original dataset is small or lacks sufficient variation to adequately represent the problem being solved. The specific data augmentation techniques used in this study are shown in Figure 2.

**Figure 2.**
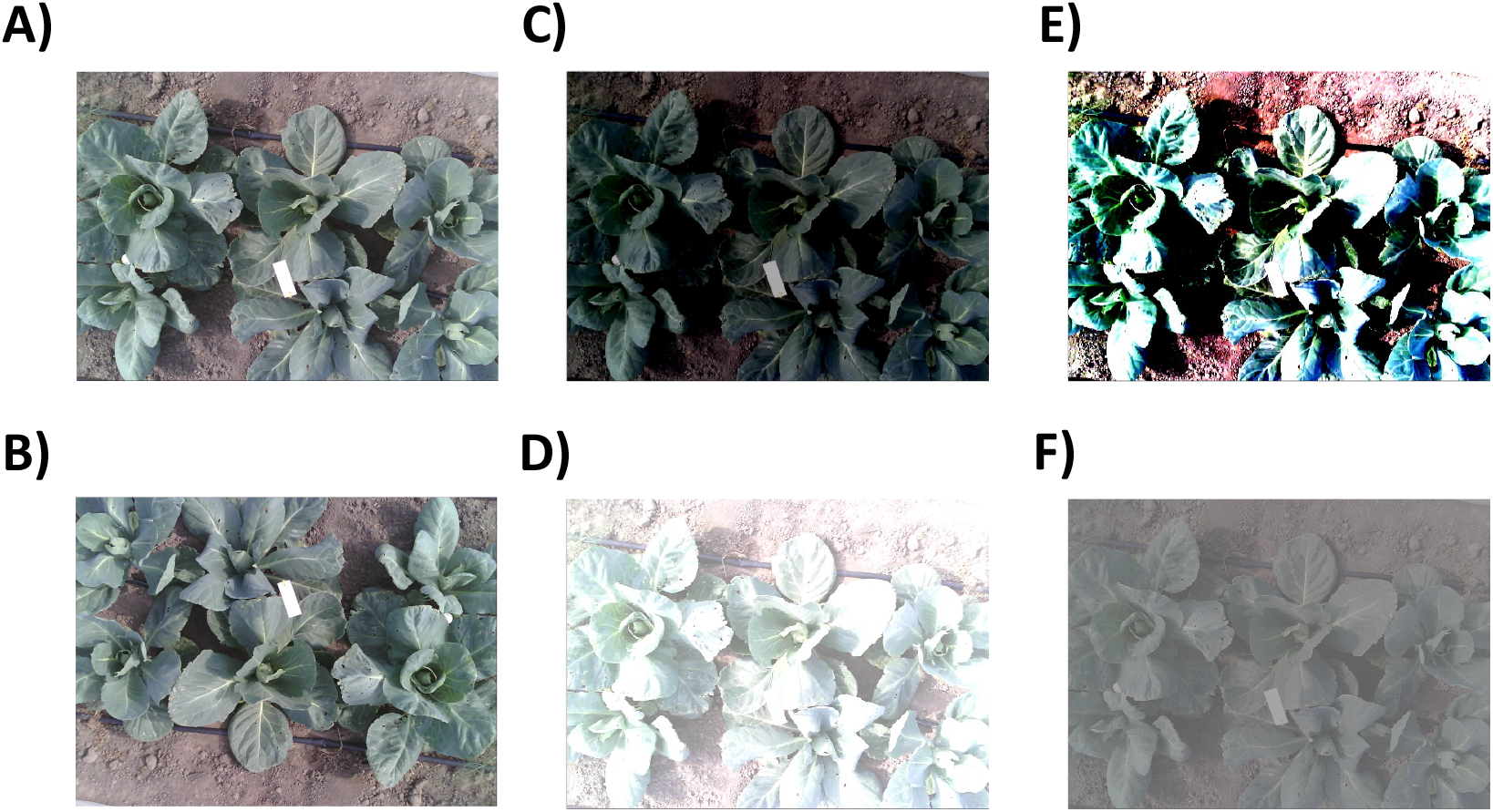
Types of data augmentation used in this study. A) Original image. B) Image rotated at a random angle (here 180°). C) and D) Decrease and increase in luminosity, respectively. E) and F) Increase and decrease in contrast, respectively.

### 2.6 Step 5: Using the trained fully convolutional neural network to perform inference on multi-species images

After training the FCNN, the next step is to use it to perform inference on multi-species images. Inference refers to the process of using the trained model to make predictions on new data. In this case, the FCNN will be used to predict the class (species) of each pixel in a multi-species image.

### 2.7 Hardware

To train the fully convolutional neural network, we utilized the Centre for High-Performance Computing and Mass Storage (CISM) at Uclouvain. CISM provides access to multiple clusters of computing units, and we used the Manneback cluster as recommended by an expert. The graphics processing unit (GPU) used was an A100-PCIE with 40 Gigabytes of memory from Nvidia. More information about the GPU can be found in the datasheet available at: https://www.nvidia.com/content/dam/enzz/Solutions/Data-Center/a100/pdf/A100-PCIE-Prduct-Brief.pdf

## 3 Results

### 3.1 Binary pseudo-labels generation

The results presented in Table 3 suggest that pseudo labeling is an effective technique for generating a training set of mono-species canopies images, particularly when done with LeafPainter or RootPainter. Furthermore, it appears that only a small amount of time ( 7 minutes) is required to achieve a high level of accuracy when using this method. Further analysis reveals that the training of a fully convolutionnal neural network on the pseudo-labels generated by Interactive Machine Learning to regenerate those pseudo-labels consistently leads to better results compared to the use of Interactive Machine Learning alone. These finding suggest that pseudo labeling can be a valuable tool in the training process, especially when combined with the training of a fully convolutional neural network. The results suggests that this combination of techniques is an effective approach in terms of producing an accurate training set at lower cost.

**Table 3:**
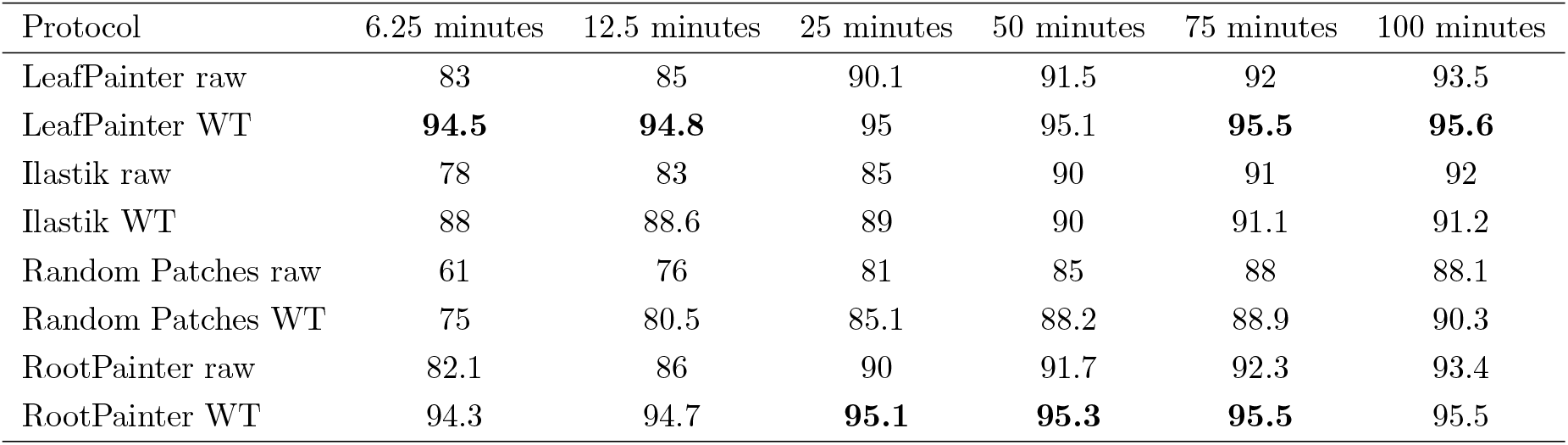
Accuracy (%) of the binary masks generated according to the protocol used and the total annotation time allocated for the given protocol.

| Protocol | 6.25 minutes | 12.5 minutes | 25 minutes | 50 minutes | 75 minutes | 100 minutes |
| --- | --- | --- | --- | --- | --- | --- |
| LeafPainter raw | 83 | 85 | 90.1 | 91.5 | 92 | 93.5 |
| LeafPainter WT | **94.5** | **94.8** | 95 | 95.1 | **95.5** | **95.6** |
| Ilastik raw | 78 | 83 | 85 | 90 | 91 | 92 |
| Ilastik WT | 88 | 88.6 | 89 | 90 | 91.1 | 91.2 |
| Random Patches raw | 61 | 76 | 81 | 85 | 88 | 88.1 |
| Random Patches WT | 75 | 80.5 | 85.1 | 88.2 | 88.9 | 90.3 |
| RootPainter raw | 82.1 | 86 | 90 | 91.7 | 92.3 | 93.4 |
| RootPainter WT | 94.3 | 94.7 | **95.1** | **95.3** | **95.5** | 95.5 |

One of the key benefits of using pseudo-labels is the ability to improve the quality of the masks. In our study, we found that the addition of the FCNN training on the IML generated pseudo-labels resulted in a significant improvement in the quality of the masks. This is illustrated on figures 3 and 4.

**Figure 3.**
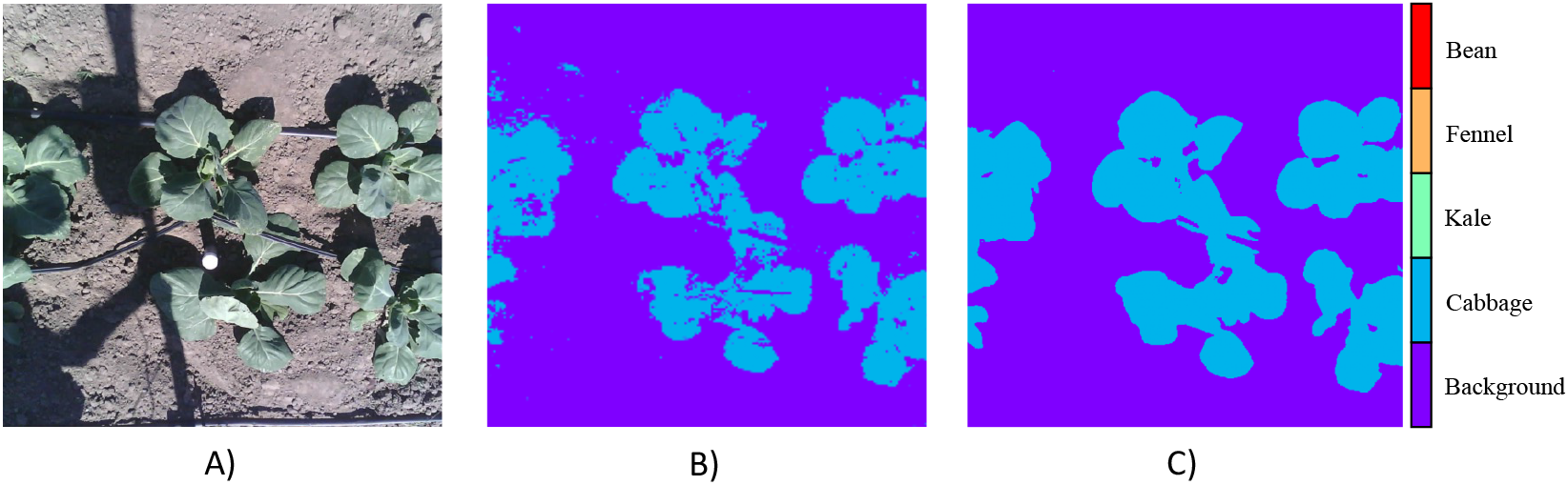
Image of a pure culture of cabbages. A) Photo of the plot. B) Binary segmentation performed by LeafPainter. We see that some areas are segmented in a sparse way. Particularly in the shadow areas. C) Generated pseudo-label. We see that the segmented zones are compact zones, which makes it possible to overcome the scattered segmentation zones

**Figure 4.**
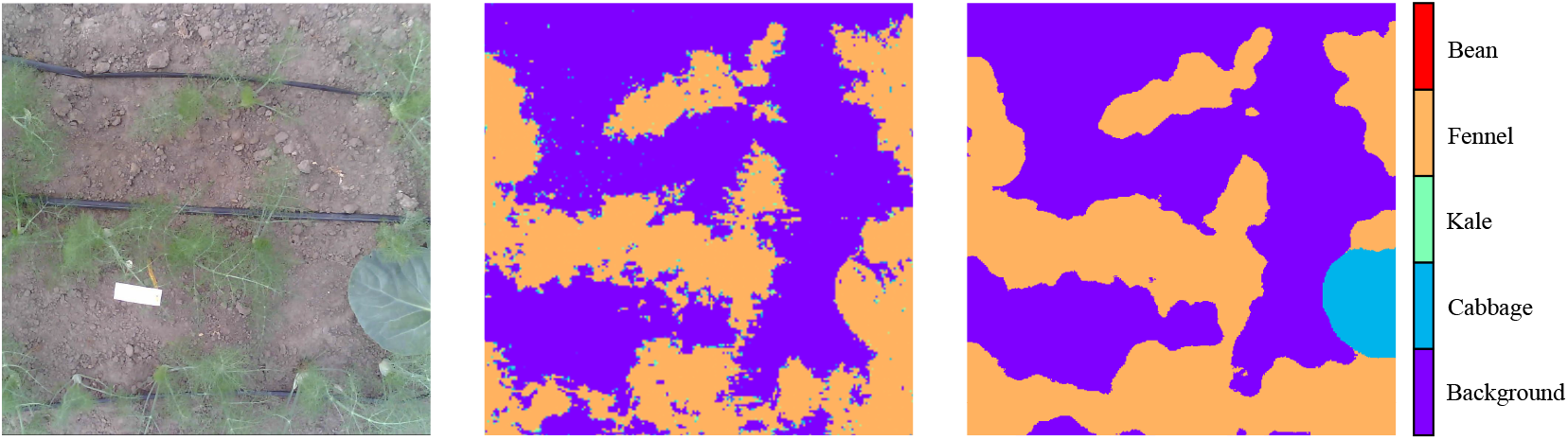
Pure culture of fennel with intrusion of cabbage. A) Image categorized as a pure crop of Fennel with head cabbage intrusion into the black circle. B) Labeling by LeafPainter. As expected, the head cabbage intrusion is labeled as fennel. C) The intrusion is correctly recognized as head cabbage by the FCNN.

### 3.2 Multiclass segmentation with the fully convolutional neural network

The segmentation is effective for various combinations of associations, as illustrated in Figure 5, even when the plants are highly intertwined.

**Figure 5.**
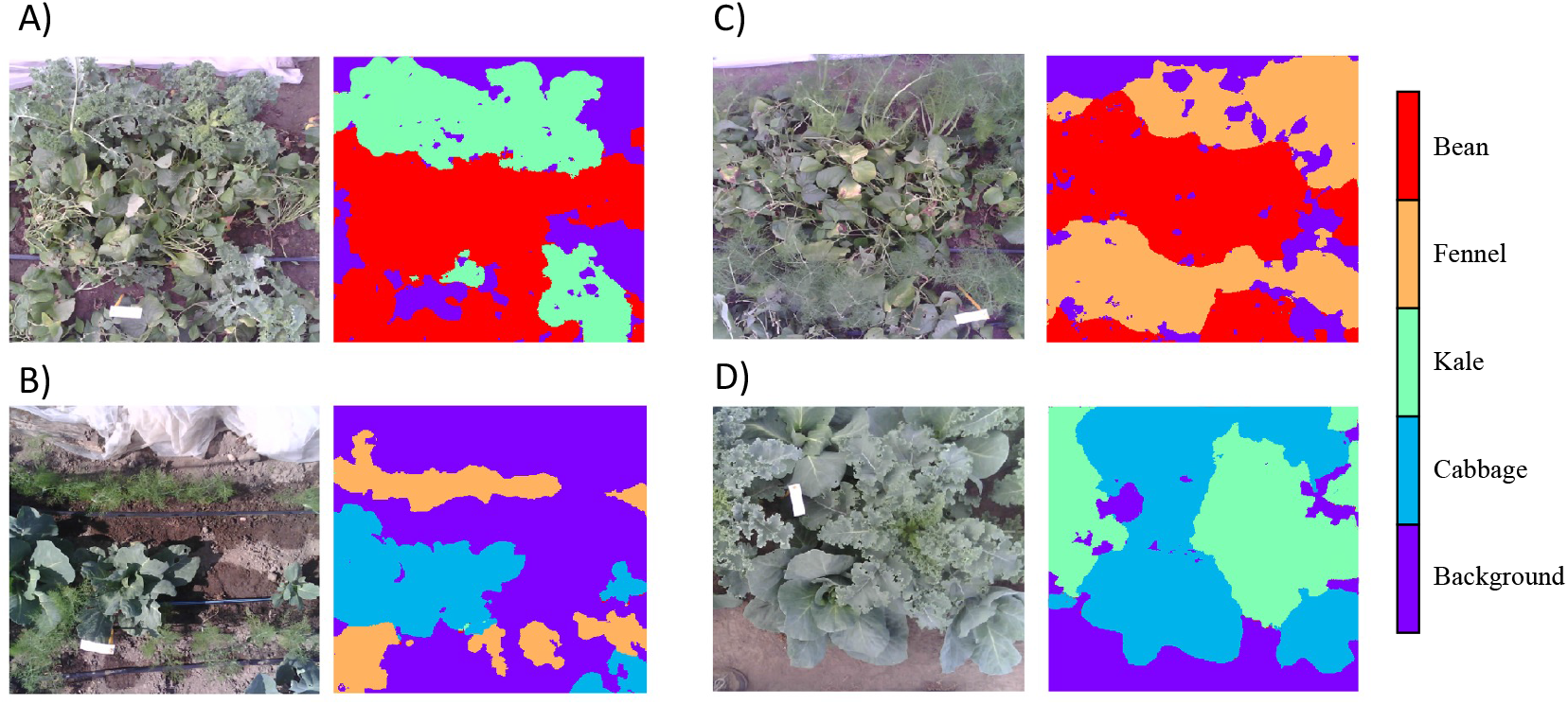
Example of different combinations of crop associations. A) Association of fennel and bean. The segmentation is well done even though the plants are very tangled. B) Association of fennel and head cabbage. Apart from a few minor errors, the segmentation is well done. C) Association of beans and fennel. The segmentation is good despite the entanglement of the plants. D) Association of head cabbage and kale. Again, the segmentation is good despite a large entanglement between the plants.

Table 4 shows that the results of the two classification tasks, binary and multiclass, are similar. The protocols that perform well for binary classification also tend to perform well for multiclass classification. However, the overall performance for multiclass classification is consistently lower than for binary classification, with the difference in performance being similar across all protocols. This likely occurs because multiclass classification is generally more challenging than binary classification.

**Table 4:** Accuracy (%) of the final multiclass masks generated according to the protocol used and the total annotation time allocated for the given protocol.

| Protocol | 6.25 minutes | 12.5 minutes | 25 minutes | 50 minutes | 75 minutes | 100 minutes |
| --- | --- | --- | --- | --- | --- | --- |
| LeafPainter raw | 78.5 | 80.6 | 85.7 | 87.1 | 87.7 | 89.1 |
| LeafPainter WT | **90** | 90.4 | **90.7** | 90.8 | **91.1** | 91.3 |
| Ilastik raw | 73.7 | 78.8 | 80.7 | 85.9 | 86.9 | 87.9 |
| Ilastik WT | 83.8 | 84.3 | 84.7 | 85.9 | 86.6 | 87.0 |
| Random Patches raw | 56.8 | 71.7 | 76.8 | 80.3 | 83.4 | 83.9 |
| Random Patches WT | 70.8 | 76.1 | 81.0 | 83.7 | 84.5 | 85.9 |
| RootPainter raw | 77.5 | 81.5 | 85.5 | 87.2 | 87.9 | 88.9 |
| RootPainter WT | 89.8 | **90.5** | 90.6 | **90.9** | **91.1** | **91.4** |

## 4 Discussion

Our research shows that monospecies images can be used to train a multispecies segmentation model. A benefit of this approach is that monospecies images are often more readily available and easier to handle than multispecies images. Additionally, training on monospecies images allows for binary segmentation, where the model only needs to distinguish between the target plant and the background. While labeling monospecies images with the species name may be required for converting binary masks to multispecies ones, this is a minor inconvenience that can be easily addressed by including the species name in the image collecting protocol. Overall, the availability and simplicity of binary segmentation offered by monospecies images make them a valuable option for training a multispecies segmentation model.

### 4.1 Binary pseudo-labels generation

In our study, we assessed different methods and tools to create an appropriate training set for multispecies canopy segmentation. Our findings indicated that RootPainter and LeafPainter were the most effective Interactive Machine Learning tools for this purpose, providing the most precise segmentation masks for the same amount of human effort. Moreover, using these tools in combination with the training of a fully convolutional neural network enhanced the quality of the segmentation masks and enabled pseudo-labels generation on the entire dataset, which made the training set much richer. Our study also revealed that the pseudo-labels generated using these methods were of superior quality compared to those produced through Interactive Machine Learning alone. Importantly, using pseudo-labels didn’t require additional human time, resulting in a more efficient and cost-effective training process. In contrast, the widely used Ilastik Interactive Machine Learning tool was surpassed by RootPainter and Leafpainter because of the slower processing speed of its random forest algorithm when compared to neural networks and gradient-boosted trees. It shows that the rate of mask creation is a crucial factor in Interactive Machine Learning and pseudo labelling as it affects the efficiency of the process. Our results highlight the importance of choosing an efficient tool and integrating pseudo-labels in the training process to achieve effective segmentation.

Annotation time is a crucial but often overlooked parameter in supervised segmentation work. It is often the most expensive part of the process. However, many authors forget to mention it in their research. For example, in Sheikh2020, different active learning methods were compared but only based on the number of training samples needed to achieve the same performance, not on the time needed to process these samples. Minimizing the amount of data needed for training is important, but the main objective should be to minimize working time. In many cases, A large, but quick-to-create, less-accurate dataset is more desirable than a light, accurate, but time-consuming dataset. This principle is demonstrated by the emergence of synthetic datasets. Sayez2022 also come to this conclusion, showing that a coarse annotation can require up to 10 times less time than a high-quality annotation, for the same performance objective. Therefore, it is crucial that future research in this area also report the time efficiency of their methods. A common dataset of crop images dedicated to the evaluation of the temporal efficiency of annotation methods would be ideal for fair comparisons. Additionally, tools like LeafPainter and RootPainter are not limited to plant segmentation and could easily be adapted to other domains for future research to explore.

### 4.2 Multi-class segmentation

Fully convolutional neural networks have demonstrated to exhibit a strong ability to generalize their learning from a single species dataset to a multi-species dataset. This has been shown to be particularly useful in the context of image segmentation tasks, where the FCNN is able to extract relevant features from the input data regardless of the specific species present. It has also been observed that there is a strong relation between the quality of the binary pseudo labels used to train the model and the overall performance of the multiclass model. Therefore, it is crucial that the quality of the pseudo labels is high in order to ensure that the model is able to effectively learn from them and improve its performance.

In this work, we have demonstrated the effectiveness of using Interactive Machine Learning and pseudo-labeling techniques to address the issue of limited human time in image segmentation. By incorporating human feedback and guidance into the training process of a machine learning algorithm, IML enables the algorithm to quickly extract meaningful features from a small set of images. Additionally, by using the model’s own predictions as labels for unlabelled data, pseudo-labeling allows the model to generalize the features it has learned to the larger dataset.

The combination of these two techniques has been shown to be an effective approach for solving image segmentation problems. Our studies have shown that good results can be achieved with just 7 minutes of human input, compared to the hours of manual annotation required by traditional supervised learning methods. This represents a significant increase in efficiency and highlights the potential of IML and pseudo-labeling for addressing image segmentation problems in cases where human time is a constraint. Our work provides evidence for the potential of these techniques to efficiently extract meaningful features and generalize them to the whole dataset.

### 4.3 LeafPainter and RootPainter

Despite having observed a similar performance between the 2 softwares, a likely advantage of LeafPainter, is that user input is directly used in inference, whereas with RootPainter we only used annotation for model training, meaning the model didn’t necessarily fit the annotations for all images, especially not the annotation in the validation set. However, RootPainter is a much more mature and tested software.

## 5 Conclusion

In this work, we focused on using machine learning techniques to improve the process of semantic segmentation for mixed crops images. Semantic segmentation involves labeling each pixel in an image with a corresponding class, such as a specific type of plant or background. This is a crucial task for studying the eco-physiological characteristics of multispecies canopies, but it can be time-consuming and labor-intensive when done manually.

To address this issue, we demonstrated that a fully convolutional neural network can generalize from single species canopy images to multispecies canopy images. This means that the FCNN is able to accurately classify pixels in images of mixed species canopies, even if it was only trained on images of single species canopies.

We also introduced Interactive Machine Learning and pseudo labeling as a way to generate single species canopy datasets much faster than traditional annotation techniques. These techniques involve using a combination of human input and machine learning algorithms to label images, which can be much more efficient than manual annotation.

We compared several methods of applying this approach and found that using LeafPainter or RootPainter for binary pseudo labels generation outperforms all the tested protocols.

Finally, being able to segment multispecies canopies at a lower cost should make it easier and faster to study the spatio-temporal dynamics of canopy area evolution, as done by Chevalier2022, partly with this method. Our method will also help future works to increase the speed of multispecies canopies segmentation and thus allow the whole scientific community to have more segmented images of crop associations and hence accelerate the progress of understanding the complex mechanisms that determine the success or failure of a crop association.

## Author Contributions

Charles Rongione conducted most of the experiments, wrote the article and implemented LeafPainter. Abraham George Smith largely contributed to the experimental design of the RootPainter evaluation and to the paper design. Céline Chevalier collected the images constituting the dataset. Xavier Draye, Guillaume Lobet and Christophe De Vleeschouwer supervised the project and reviewed the writing of the paper.

